# Enabling MHC-mismatched hematopoietic stem cell transplants and organ graft tolerance without chemotherapy or radiation

**DOI:** 10.1101/525899

**Authors:** Benson M. George, Kevin S. Kao, Angela Chen, Alan C. Le, Akanksha Chhabra, Cassandra Burnett, Kyle M. Loh, Judith A. Shizuru, Irving L. Weissman

## Abstract

Hematopoietic stem cell (HSC) transplantation can replace diseased blood systems with a healthy one, thereby treating or curing genetic blood and immune disorders including autoimmune diseases and immunodeficiencies. However, toxic chemotherapy or radiation is necessary to ablate an animal’s existing blood system prior to HSC transplantation, leading to significant morbidity. To accomplish safer blood-system replacement we developed a combination of six monoclonal antibodies to safely and specifically deplete the HSCs, T cells and NK cells of immune-competent mice. Remarkably, immunologically-foreign (allogeneic) HSCs mismatched at half or all the *MHC* genes could engraft these antibody-treated mice, generating donor blood systems that stably co-existed with host blood cells. These chimeric immune systems were immunologically tolerant to heart tissue from the HSC donor, providing a safe platform for HSC transplantation as a means to solid organ transplantation. The ability to transplant *MHC*-mismatched HSCs without chemotherapy or radiation has significant ramifications for regenerative medicine.

## Introduction

A multitude of genetic blood and immune system disorders can be treated by hematopoietic cell transplantation (HCT): examples include thalassemia, sickle cell anemia, Fanconi’s anemia, inherited immunodeficiencies, autoimmune diseases (e.g., type 1 diabetes), and metabolic storage disorders^8-13^. These diseases can be corrected when an individual’s blood system is replaced by healthy, transplanted blood cells, which stably derive from the rare hematopoietic stem cells (HSCs) in HCT grafts^14,15^. After regeneration of a donor-derived blood and immune system, HCT recipients are immunologically tolerant to organ transplants from the HSC donor^16-18^. While any single-gene or multi-gene genetic disorder of the blood system could be cured by allogeneic HCT, treatment of non-malignant hematological or immunological disorders only accounted for 6% of total HCT cases reported in Europe in 2015^19^.

To overcome the disproportionately infrequent use of HCT to treat non-malignant blood disorders and extend its reach, two key challenges must be addressed: safety concerns and donor availability. At present, allogeneic HCT leads to clinical or subclinical graft vs. host disease (GvHD) caused by contaminating donor-derived T cells^20,21^; GvHD can be overcome by transplanting purified HSCs devoid of T cells. Moreover, HCT conditioning requires chemotherapy and radiation, which can induce life-threatening side effects^22,23^. Another challenge confronting HCT for genetic blood disorders is the current need for fully matched donors at the human leukocyte antigens (HLA, otherwise known as major histocompatibility complex [MHC]) loci; while 75% of Caucasian Americans currently have matched donors, it is markedly harder to find fully-matched donors for Black Americans (16-19% currently have a match) or other under-represented ethnic groups^24^. If it were possible to safely perform HCT using haploidentical donors (which are matched at half of HLA loci), this would significantly expand the availability of donors to theoretically enable any individual to receive HCT from their parent, child, or 75% of siblings. Although some monogenic hematological disorders can be cured with gene-editing, diseases with multiple genetic changes cannot at this time be approached by gene editing, but can be approached with healthy HSCs. In addition, healthy donor HSCs can block a number of autoimmune diseases, and tissue and organ grafts from HSC donors are tolerated in allogeneic hosts. Therefore, if it were possible to safely transplant fully HLA-mismatched HSCs, this would massively open the pool of available donors and patients treatable with HSC transplants; moreover, recipients would be immunologically tolerant to foreign organs or tissues obtained from the same donor. This could enable HLA-mismatched organ transplants without the lifelong immunosuppression commonly needed to prevent rejection of vital organs^25^.

The safety of HCT could be considerably improved if toxic conditioning regimens (chemotherapy and/or radiation) could be partially or wholly replaced by more specific agents, such as monoclonal antibodies. However, many previous studies attempting this still relied upon low dose radiation/chemotherapy or unfractionated bone marrow likely contaminated with GvHD-inducing T cells^26-29^, the effects of which can be overt or subclinical^30^. While extant antibody conditioning regimens enable the transplantation of minor histocompatibility antigen-mismatched HSCs^31-33^, it has not been possible to transplant MHC-mismatched HSCs using antibody-based conditioning. Here we demonstrate that conditioning using six monoclonal antibodies enables wild-type mice to receive partially-(haploidentical) or fully-MHC mismatched HSCs, therefore enabling blood system replacement and induction of tolerance to mismatched donor organs without recourse to chemotherapy or radiation.

## Results

For haploidentical transplantation experiments, AKR x C57BL/6 F_1_ (hereafter referred to as AB6F_1_) mice were used as bone marrow or HSC donors and BALB/C X C57BL/6 F_1_ (CB6F_1_) (**Fig. 1a**) mice served as recipients; these mouse strains are matched at the *H2*^*b*^ haplotype but mismatched for *H2*^*k*^ and *H2*^*d*^ (i.e., at half of the Major Histocompatibility Complex [MHC] haplotypes) (**Fig. 1b**). We sought to determine if conventional conditioning could be replaced with monoclonal antibodies (mAb). We previously demonstrated that immune-deficient mice could be conditioned using an anti-c-Kit antibody to enable syngeneic HSC engraftment^32^, whereas comparable conditioning of immune-competent mice required dual administration of anti-c-Kit and anti-CD47 blocking agents^33^. CD47 blockade enables macrophages to phagocytose antibody-bound (opsonized) cells^34,35^, such as c-Kit^+^ HSCs opsonized by anti-c-Kit antibodies.

**Figure 1.**
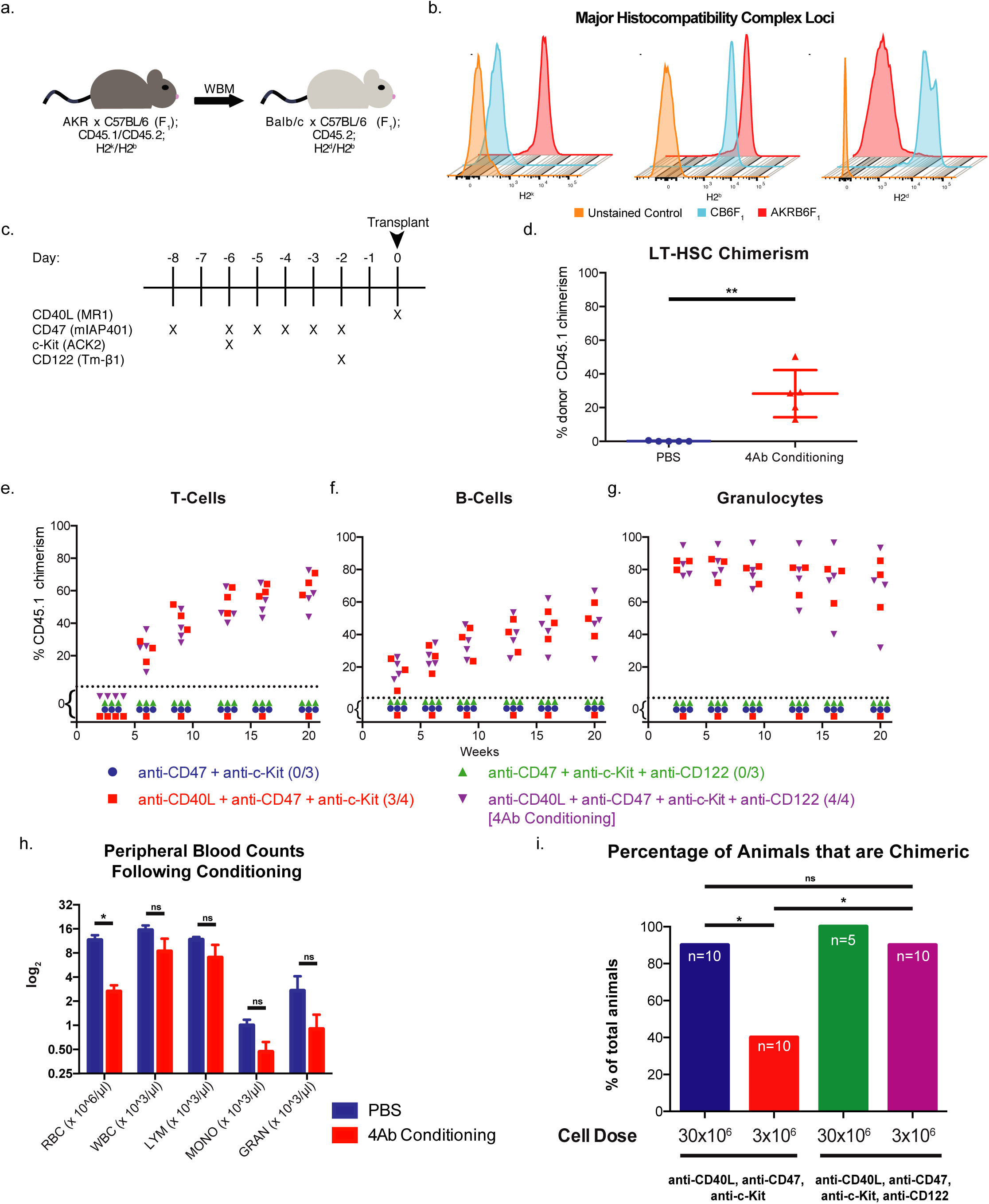
A monoclonal antibody cocktail can induce long-term multi-lineage hematopoietic reconstitution. (a) Haploidentical transplantation schema using AKRB6F_1_ donors and CB6F_1_ recipients. (b) Flow cytometric analysis of MHC Class I on donor and recipient strains. (c) Dosing schedule for 4Ab conditioning regimen. (d) Twenty-four-week donor chimerism in the long-term HSC compartment (Lin^-^ c-Kit^+^ Sca1^+^ CD150^+^ Flk2^-^ CD34^-^) following PBS or Ab conditioning (n=5). (e-g) Chimerism over time following variations of 4Ab conditioning and WBM transplantation (n=3-4). (h) CBC following 4Ab conditioning on Day 0 (n=3). (i) Percentage of animals which are chimeric at 3 weeks after various WBM doses, with or without NK cell depletion (n=5-10). Data and error bars in (d) and (h) represent means ± SD; (d) unpaired t-test reveals P≤0.01 and multiple t-test reveals P≤0.05. Data in (i) represent means; one way ANOVA was performed where *P≤0.05.

However, in order to engraft allogeneic HSCs mismatched at the MHC loci, we hypothesized that it is crucial to suppress or eliminate both T cells and NK cells, which reject cells expressing foreign, minor histocompatibility antigens^36,37^ or that lack “self” MHC^38^. To eliminate host NK cells we targeted CD122/Il2Rβ (which is expressed throughout human and mouse NK cell development^39^) using the anti-CD122 mAb Tm-β1^40,41^. To prevent T-cell mediated rejection we targeted CD40L (also known as CD154), which is a co-stimulatory cell surface molecule expressed by activated T-cells and is required for their signaling with CD40^+^ antigen presenting cells^42^. Interruption of the CD40-CD40L axis can help induce tolerance to hematopoietic cells and skin grafts^43-45^ and importantly, does not deplete all T cells since CD40L is upregulated on activated T cells; we inhibited CD40L using the anti-CD40L antibody MR1^46^.

Mice were treated over the course of nine days (**Fig. 1c**) with the four monoclonal antibodies (anti-CD122, anti-CD40L, anti-c-Kit and anti-CD47; herein referred to as 4Ab conditioning) and then transplanted with 30 million whole bone marrow (WBM) cells. Chimerism, which is the presence of >1% donor cells in peripheral blood lineages (although higher percentages are preferred to ameliorate pathological situations), was periodically measured by CD45 allelic differences (**Extended Data Fig. 1a**) and multi-lineage mixed chimerism was observed in all animals receiving 4Ab conditioning (**Extended Data Fig. 1b-d**). Importantly, mixed chimerism was also observed in the long-term HSC (LT-HSC) compartment (**Fig. 1d**), indicating that the donor chimerism did not result from engraftment of long-lived mature immune cells, but was being actively maintained by donor stem cells.

To identify the minimally necessary components of this cocktail, we tested each antibody in isolation (**Extended Data Fig. 2**) and in various permutations. The minimally necessary cocktail to engraft 30 million WBM cells was anti-CD47, anti-c-Kit, and anti-CD40L (**Fig. 1e-g**). However, only 75% of the mice in the group lacking anti-CD122 were chimeric, while the complete 4Ab conditioning induced chimerism for all mice. Interestingly, engrafted animals from both groups showed similar levels of multi-lineage chimerism over twenty weeks. Additionally, the 4Ab conditioning did not induce granulocytopenia prior to transplantation (**Fig. 1h**). We tested the lowest dose of WBM that could engraft by titrating the dose of WBM while modulating the usage of anti-CD122. The amount of peripheral blood chimerism decreased as the amount of bone marrow transplanted decreased (**Extended Data Fig. 3**). When anti-CD122 was excluded from 4Ab conditioning, the percentage of mice that became chimeric significantly decreased with a decreasing amount of bone marrow. However, anti-CD122 enabled engraftment in a similar percentage of mice despite decreases in bone marrow dose (**Fig. 1i**).

To eliminate the possibility of GvHD, next we transplanted enriched HSC populations (as opposed to WBM). In these experiments Lineage^-^ Sca1^+^ c-Kit^+^ (LSK) cells (**Fig. 2a**) were transplanted, which are highly enriched for HSC and multipotent progenitor (MPP) cells^47,48^. Both c-Kit-enriched and LSK cells were given in quantities that corresponded to their abundance in 3 million WBM cells (**Fig 2b**). All three types of grafts showed complete, long-term multi-lineage chimerism in irradiated controls. Strikingly, while 4Ab-conditioned mice were successfully engrafted long term by WBM, they were not reconstituted by c-Kit-enriched or LSK transplants (**Fig. 2b**). This therefore indicates that additional conditioning antibodies are required for enriched HSC populations to successfully engraft.

**Figure 2.**
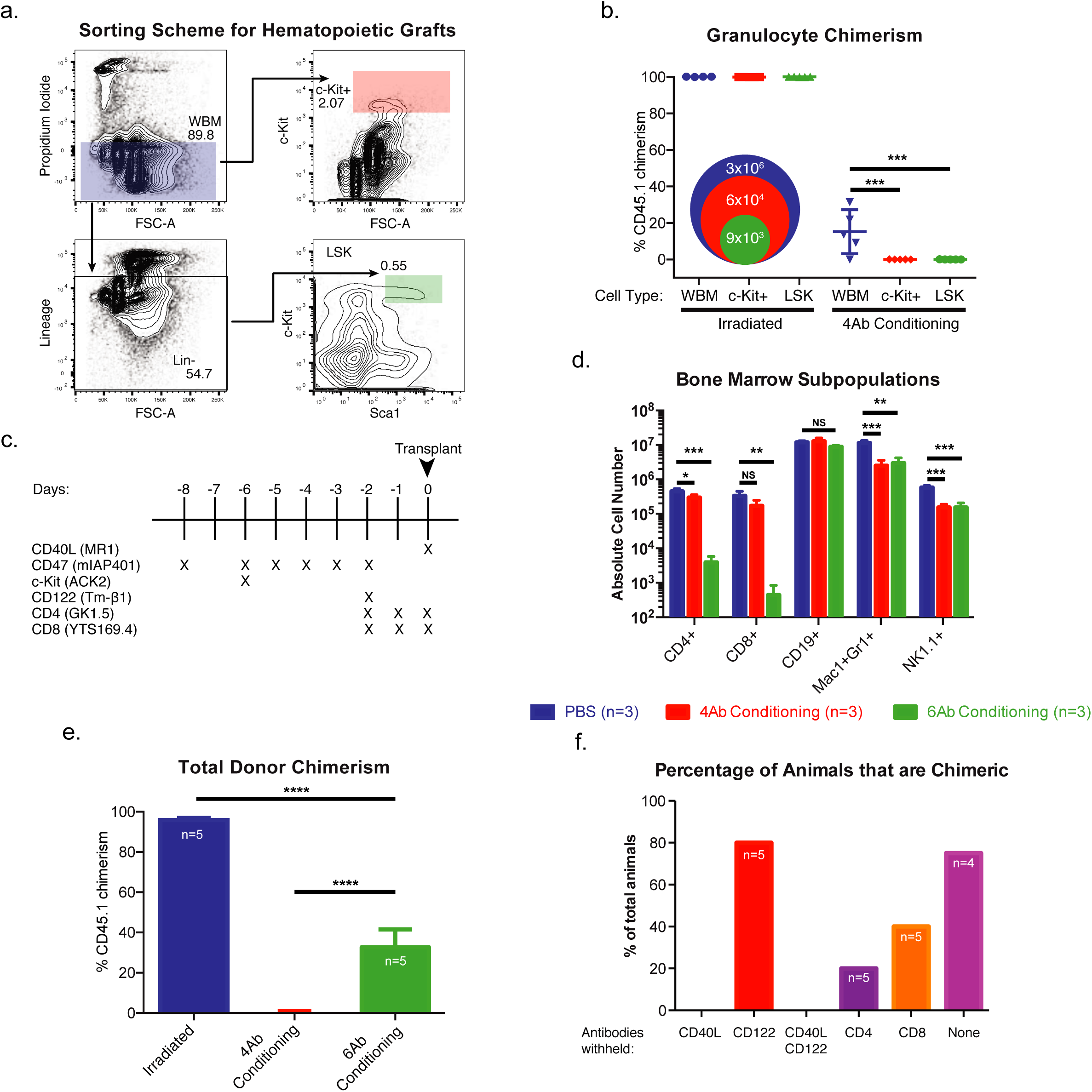
A monoclonal antibody cocktail can induce long-term multi-lineage hematopoietic reconstitution of low dose purified HSCs. (a) Sorting scheme used to calculate and isolate LSK and c-Kit^+^ cells for transplantation with representative percentages of cell abundance. (b) Granulocyte chimerism at 16 weeks following WBM, Kit+, and LSK transplantation with 4Ab conditioning or irradiation; 9×10^3^, 6×10^5^, and 3×10^6^ cells were given for LSK, Kit+, and WBM, respectively (n=4-5). (c) Dosing schedule for 6Ab conditioning. (d) Mature immune cell population abundances in the bone marrow following 6Ab conditioning (n=3). (e) Total peripheral blood donor chimerism following LSK transplantation at 16 weeks (n=5). (f) Percentage of animals which are chimeric following exclusion of individual components of the 6Ab cocktail (n=4-5). Data and error bars in (b), (d) and (e) represent means ± SD; (b-e) one way ANOVA was performed where *P≤0.05, **P≤0.01, and ***P≤0.001.

In order to facilitate LSK engraftment we attempted to provide additional immune suppression by eliminating T cells using anti-CD4 and anti-CD8 depleting antibodies^33,49^ (**Fig. 2c**). The addition of anti-CD4 and anti-CD8 antibodies to the 4Ab regimen robustly depleted T-cells from peripheral blood, spleen and bone marrow (**Fig. 2d** and **Extended Data Fig. 4**). The usage of this six antibody cocktail, which will be referred to as 6Ab conditioning (anti-CD122, anti-CD40L, anti-c-Kit, anti-CD47, anti-CD4 and anti-CD8 mAbs), induced multi-lineage long term chimerism in recipients transplanted with 9000 LSK cells (**Fig. 2e and Extended Data Fig. 5a-c**). Additionally, due to this lymphoablative approach, we monitored T-cell kinetics in this cohort and so that peripheral T-cells were present 3-weeks post-transplant and continued to rise until week 8 (**Extended Data Fig. 5d**). Lastly, the total LSK cell dosage corresponds to approximately 360,000 LSK/kg, which is well below HSC doses seen in preclinical testing for allografts in mice^50^ and clinical usage in autografts in humans^51^. In summary, 6Ab conditioning enables low doses of HSC to engraft mice without recourse to chemotherapy or radiation.

It was unclear if all six components of this cocktail were necessary; therefore, we used a reductive process to identify the dispensable antibodies. Removal of anti-CD40L, anti-CD4, and anti-CD8 resulted in fewer chimeric animals and lower chimerism within each cohort, as compared to the complete 6Ab conditioning cohort (**Fig. 2f** and **Extended Data Fig. 6**). However, removal of the anti-CD122 antibody did not significantly change the percentage of chimeric animals as compared to the control cohort. Unlike in the 4Ab conditioning regimen, CD122 may be less necessary in the 6Ab conditioning. We hypothesize that this is due to NK dependence on T-cell activation^52^, which may be lost in the 6Ab conditioning regimen, as there is near complete depletion of T-cells.

Importantly, 6Ab conditioning followed by HSC transplantation induced centrally-mediated immunological tolerance to the donor genetic strain. Central tolerance implies thymic re-education of the host immune system to permit donor cell engraftment. To gauge central tolerance in these animals, we measured the presence of the V beta 6 (Vb6) TCR chain in peripheral blood. The Vb6 chain is reactive to the Mtv-7 provirus-encoding super-antigen, which is present in the AKR background^53^. Therefore, for AB6F_1_ HSCs to coexist in CB6F_1_, the CB6F_1_ endogenous Vb6^+^ T-cells must be clonally deleted. In both WBM- and LSK-transplanted animals, chimeric animals showed deletion of host Vb6^+^ T-cells (**Fig. 3a-b**). Interestingly, in the WBM cohort conditioned with anti-c-Kit, anti-CD47, and anti-CD40L, the only animal with a normal Vb6^+^ T-cell frequency also never achieved chimerism (**Fig. 3a**).

**Figure 3.**
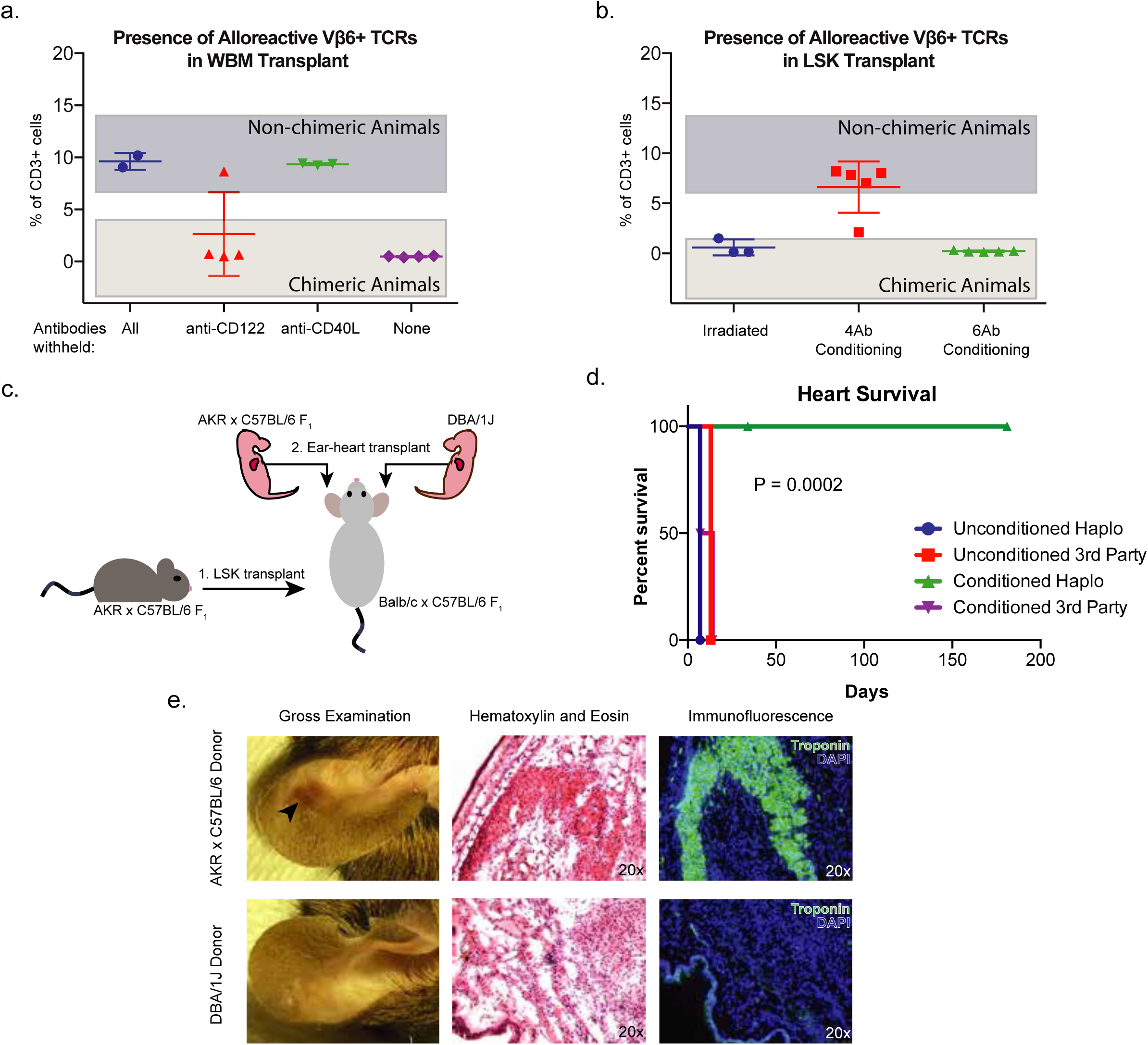
Low dose LSK transplantation via a non-genotoxic conditioning regimen allows for tolerance to donor tissue. (a-b) Abundance of donor-reactive host T-cells in peripheral blood at 22 weeks following WBM (a) and LSK (b) transplantation (n=2-5). (c) Fetal heart into ear transplantation schematic. (d) Kaplan-Meier curve showing donor heart survival (n=5). (e) Gross examination, H&E, and IF of representative ear-heart grafts 34 days following tissue transplant. Data in (d) was subjected to a log-rank (Mantel-Cox) test and yielded P = 0.0002.

Strikingly, we found that 6Ab-conditioned mice engrafted with MHC-mismatched donor HSC were immunologically tolerant to organs from the same donor strain. To this end, we transplanted heart grafts from HSC-donor (AB6F_1_) or third-party (DBA/1J strain, which are homozygous for H2^q^) newborn pups into the ear pinna of naïve and LSK-Ab conditioned chimeric animals (**Fig. 3c**). In naïve, unconditioned, untransplanted mice, both AB6F_1_ and DBA1/J hearts were rejected rapidly (**Fig. 3d**). In 6Ab conditioned chimeric mice, DBA1/J hearts were rejected within 14 days while active, beating AB6F_1_ hearts persisted for at least 181 days (**Extended Data Video 1**). Representative ear-heart grafts were harvested at 34 days and analyzed by immunohistochemistry. Upon gross examination, AB6F_1_ hearts are visible in the pinna while DBA/1J hearts are no longer apparent (**Fig. 3e**). H&E and immunofluorescence analysis showed troponin^+^ cardiac tissue lacking immune cell infiltrates in the AB6F_1_-engrafted pinna; however, by this time there was no cardiac or troponin^+^ tissue within the pinna containing DBA/1J hearts (**Fig. 3e**). This therefore indicates that MHC-mismatched donor HSC can induce immunological tolerance of 6Ab-conditioned mice to heart grafts from the same genetic donor.

Finally, we demonstrated that the 6Ab conditioning regimen enabled successful engraftment of *fully* MHC-mismatched HSCs. We used DBA1/J (H2^q^) mice as donors and CB6F_1_ (H2^b/d^) as hosts (**Fig. 4a**). After transplanting 9000 DBA1/J LSK cells, we observed high donor-host chimerism by 8 weeks in all 6Ab-conditioned CB6F_1_ mice (**Fig. 4b**). Mice transplanted with 3 million WBM cells alone failed to establish donor chimerism (confirming the necessity for conditioning), while 40% of 4Ab-conditioned mice receiving WBM achieved low levels of chimerism. Eighty-percent of the irradiated CB6F_1_ mice transplanted with WBM were dead by 9 weeks following transplantation (**Fig. 4c**), likely by GvHD, which was not observed in LSK transplants.

**Figure 4.**
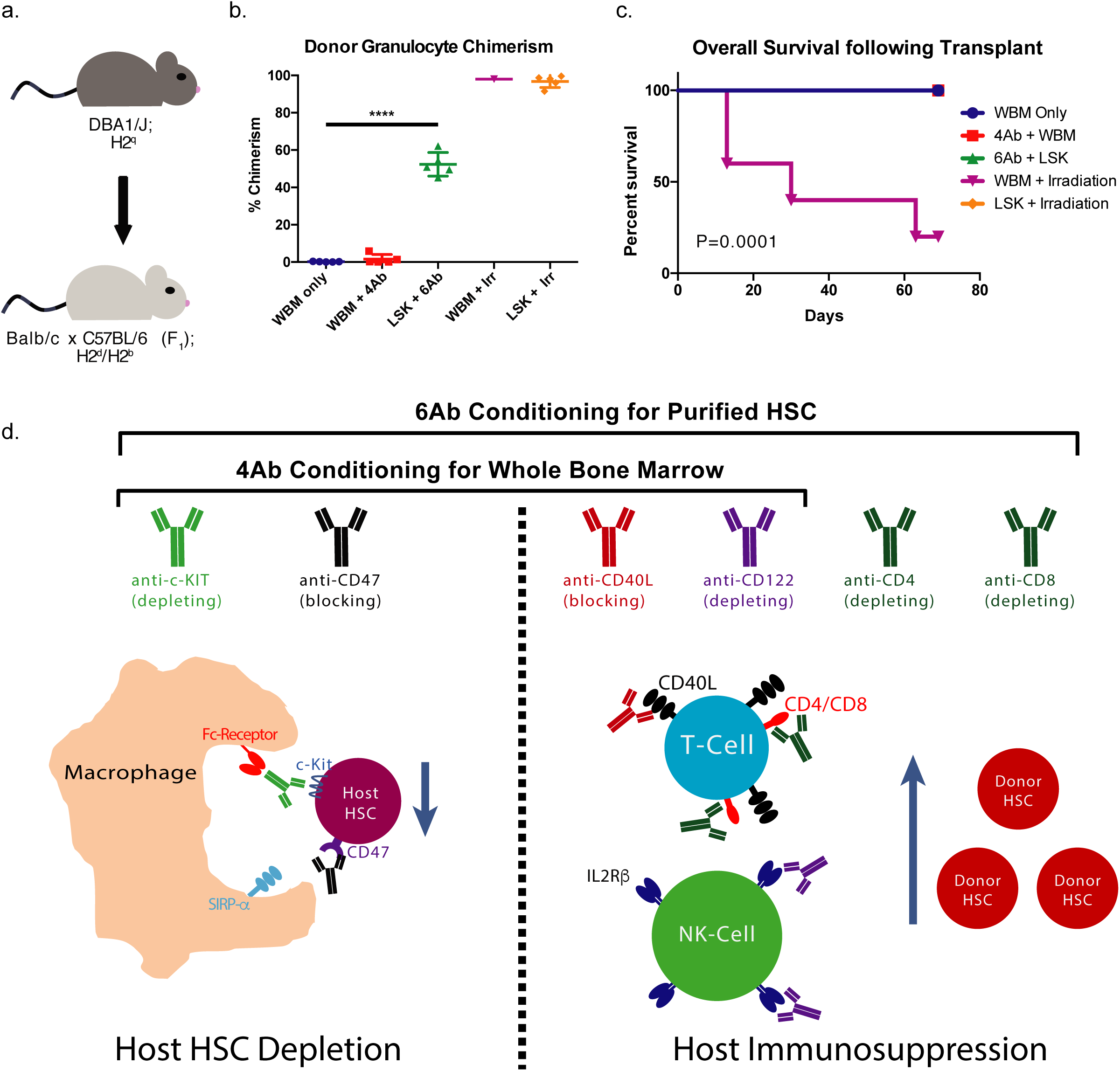
Hematopoietic stem cells can be engrafted despite a full MHC mismatch. (a) Transplantation schematic where DBA1/J are the donor and CB6F_1_ are the host. (b) Percent of donor engraftment at 8 weeks following WBM and LSK transplantation (n=5). (c) Overall survival of transplanted animals. (d) Overview describing an all-antibody conditioning regimen which can deplete endogenous HSCs, and provide transient immune suppression by targeting host T and NK cells. Data and error bars in (b) represent means ± SD and one way ANOVA was performed where ****P≤0.0001; (c) log-rank (Mantel-Cox) test yielded P = 0.0001.

## Discussion

Here we have developed a method to transplant half-(haploidentical) and fully-MHC mismatched purified HSCs into immune-competent animals; importantly this is accomplished without the use of chemotherapy and/or radiation, and without the GvHD that occurs in most, if not all, other types of HCT transplants. For instance, patients with Fanconi’s anemia are highly susceptible to DNA damage, and therefore, conventional transplant conditioning regimens pose a serious risk to this cohort. In addition, perinatal and aged patients are often excluded from allogeneic hematopoietic cell transplants, due to their susceptibility to dangerous side effects of radiation, chemotherapy, and GvHD.

This proof-of-principle study represents a potential bridge for treating non-monogenic hematological and immunological diseases where transplantation is curative but underutilized. If translatable to human patients, this has many ramifications for the treatment of blood and immune system disorders. First, this antibody conditioning regimen—combined with purified HSC transplants—could improve the safety of blood and immune system replacement by obviating the use of chemotherapy/radiation and by eliminating GvHD. Second, by facilitating transplantation of haploidentical, HLA-mismatched HSCs, this will open up the donor pool to enable most recipients to find a match, even if their age or clinical status had prevented HCT under previous protocols.

Lastly, the ability to induce immunological tolerance to foreign organs could increase opportunities for all patients requiring lifesaving organ transplants: specifically, it could obviate the need for lifelong immune suppression for patients to receive foreign organ transplants^54,55^. In particular, the immune systems in antibody-conditioned, donor HSC-transplanted animals are tolerant to donor (but not third-party) hearts. Today the donor of an organ, tissue or HSC transplant is a living or recently deceased person. The ultimate goal of regenerative medicine will be to differentiate a pluripotent (embryonic or induced pluripotent) stem cell line into HSCs and other needed tissue stem cells (such as those of the neural^56^, bone and cartilage^57^, or liver^58^), either *in vitro*^59,60^ or *in vivo* within a large-animal host (such as a pig)^61,62^. This would relieve the need for human beings to give up their HSCs and organs for others. Antibody conditioning, followed by co-transplantation of pluripotent stem cell-derived HSC and tissue stem cells, could deliver lifesaving organs for patients without recourse to long-term immunosuppression.

## Methods

### Animals

All experiments were performed according to guidelines established by the Stanford University Administrative Panel on Laboratory Animal Care. AKR x C57BL/6 F_1_ donors were crossed and bred in house. CB6F_1_ and DBA1/J recipients were purchased from the Jackson Laboratory. DBA1/J pregnant females were purchased from Taconic Biosciences for ear heart grafts.

### Antibodies

Anti-CD47 (mIAP410), anti-c-Kit (ACK2), anti-CD122 (Tm-β1), anti-CD40L (MR1), anti-CD4 (GK1.5), and anti-CD8 (YTS169.4) were purchased from BioXCell. Anti-CD47 was given intraperitoneally as a 100ug dose on Day −8 and then as a 500ug dose for subsequent injections throughout the conditioning process^33^. Retro-orbital anti-c-Kit and intraperitoneal anti-CD40L were both given as one time 500ug boluses^44^. Anti-CD122 was given intraperitoneally as a 250ug dose while anti-CD4 and anti-CD8 were given as 100ug intraperitoneal doses^41^. Mice receiving anti-c-Kit antibody were given 400ug of diphenhydramine intraperitoneally 15 minutes prior to injection.

### Graft Preparation and Transplantation

Whole bone marrow was extracted from donor mice tibia, femurs, hips, and spine. Bones were crushed, filtered, and subsequently underwent red blood cell (RBC) lysis. For c-Kit-enriched transplants, RBC lysed whole bone marrow were bound to Miltenyi CD117 MicroBeads as per the manufacturer’s instructions and collected after magnetic separation. For LSK cell transplants, RBC lysed whole bone marrow were bound to the Miltenyi Lineage Cell Depletion kit cocktail as per the manufacturer’s instructions. Flow through from the magnetic separation columns was collected and stained in PBS with 2% FBS with optimal concentrations of the following antibodies: CD3 PE (17A2), CD4 PE (GK1.5), CD5 PE (53-7.3), CD8a PE (53-6.7), B220 PE (RA3-6B2), Gr-1 PE (RB6-8C5), Mac-1 PE (M1/70), Ter119 PE (TER119), SCA1 Pe-Cy7 (D7), and CD117 APC (2B8). Propidium iodide was added as a viability stain just prior to sorting on a BD Aria. All cells for transplant were resuspended at the desired concentration in PBS with 2% FBS. Irradiation control mice were lethally irradiated with two doses of 6.5Gy prior to transplantation. All mice were anesthetized using isoflurane and then transplanted with 100uL of cell suspension via retroorbital injection.

### Peripheral Blood Chimerism

Mice were periodically bled via retroorbital bleeding into EDTA coated tubes. Blood was then incubated in 1% dextran with 5mM EDTA at 37C for 1 hour. The supernatant from each tube was extracted, lysed and then stained with optimal concentrations of the following antibodies: CD3 APC (17A2), CD19 PE-Cy7 (ebio103), Gr-1 BV421 (RB6-8C5), Mac-1 APC-Cy7 (M1/70), CD45.1 FITC (A20), and CD45.2 PE (104). Samples were analyzed on a BD Fortessa and donor versus host chimerism was distinguished based on CD45 allelic differences.

### Ear-Heart Graft

Neonatal mice were euthanized 1-2 days after birth and their hearts were harvested. Recipient mice were prepared by making a small incision on the dorsal side of their ear near the skull. Afterward, using a trocar, a pouch was created by tunneling from the incision site to the tip of the pinna. Neonatal hearts were delivered at the distal end of the pouch with the trocar. The tunnel was closed by gently pushing the lifted skin back to the dermis. Heart viability was monitored for beating by visualizing the graft through a dissecting microscope.

## Acknowledgements

We thank Agnieszka Czechowicz, Deepta Bhattacharya, and Daniel Kraft for helping to initiate this field while in the laboratory, and for discussion of the subject matter that resulted in the paper. We thank Kenneth I. Weinberg, Ryan A. Flynn, and Adam J. Rubin for editorial comments. We thank Aaron McCarty, Theresa Storm, Teja Naik, Steve Jungers, and Jessica Poyser for technical assistance. We thank Pauline Chu for providing histological services. This study was supported by the California Institute for Regenerative Medicine (RT3-07683, to I.L.W. and J.A.S.), the Stanford-UC Berkeley Stem Cell Institute, anonymous donors and the NIH Director’s Early Independence Award DP5OD024558 (to K.M.L.), and PHS Grant Number CA09302, awarded by the National Cancer Institute, DHHS (to B.M.G.).

## Author Contributions

B.M.G. wrote the manuscript. B.M.G. and A.Chh. designed experiments. B.M.G. and K.S.K. performed hematopoietic cell transplants. B.M.G., A.Che., and K.M.L. performed and analyzed heart graft studies. A.C.L. and C.B. performed data analysis. K.M.L. edited the manuscript. I.L.W. and J.A.S. supervised the project, conceived experiments and edited the manuscript.

## Statistics Note

Unpaired t-tests, one-way Analysis of variance (ANOVA), two-way ANOVA, and log-rank tests were performed to distinguish statistical differences between groups when necessary.

**Extended Data Figure 1.**
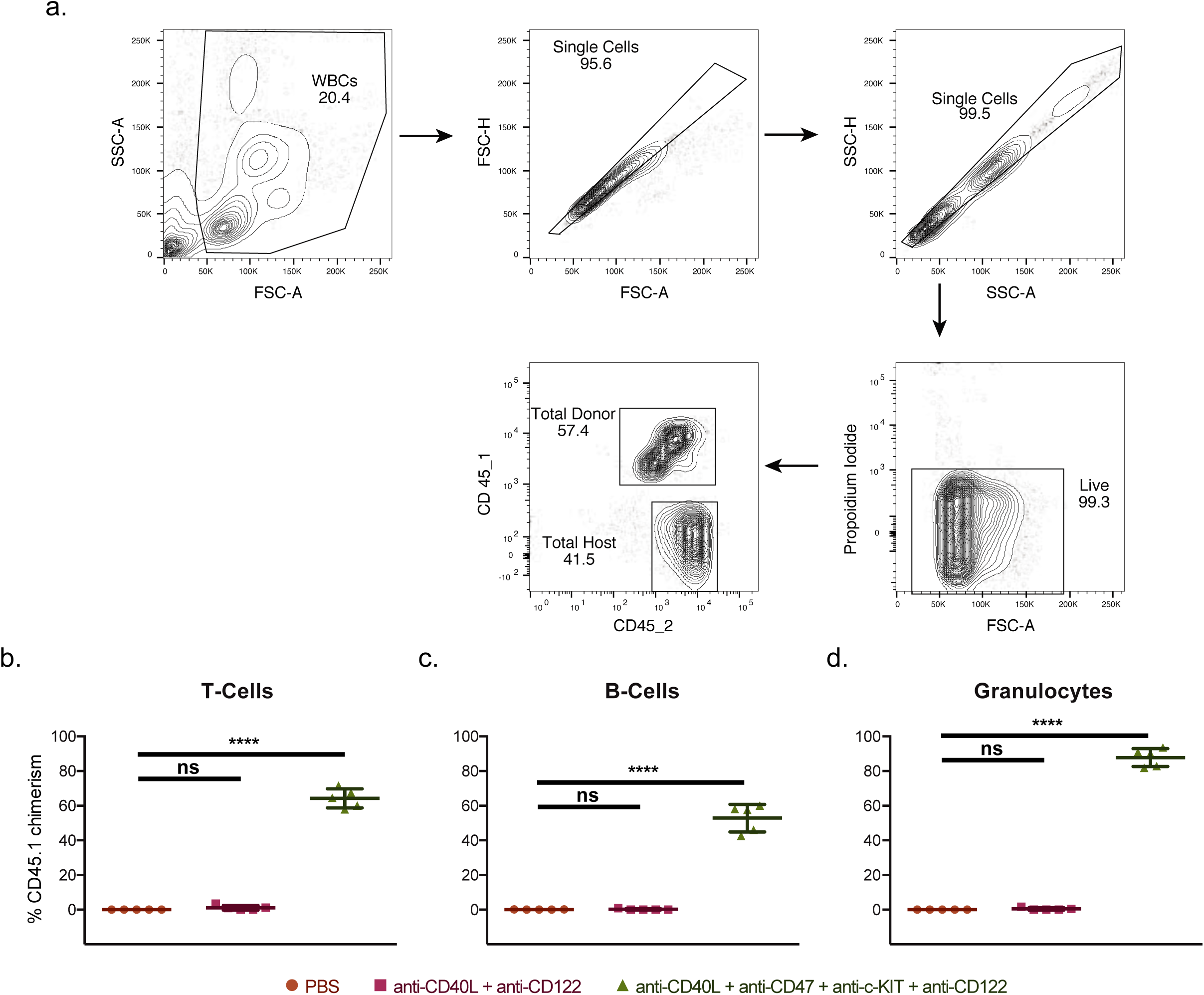
(a) FACS analysis scheme to determine peripheral blood chimerism by CD45 allelic differences between the host and the donor. (b) Multi-lineage peripheral blood chimerism 16 weeks following WBM transplant (n=5). Data and error bars in (b) represent means ± SD; one way ANOVA was performed where ****P≤0.0001.

**Extended Data Figure 2.**
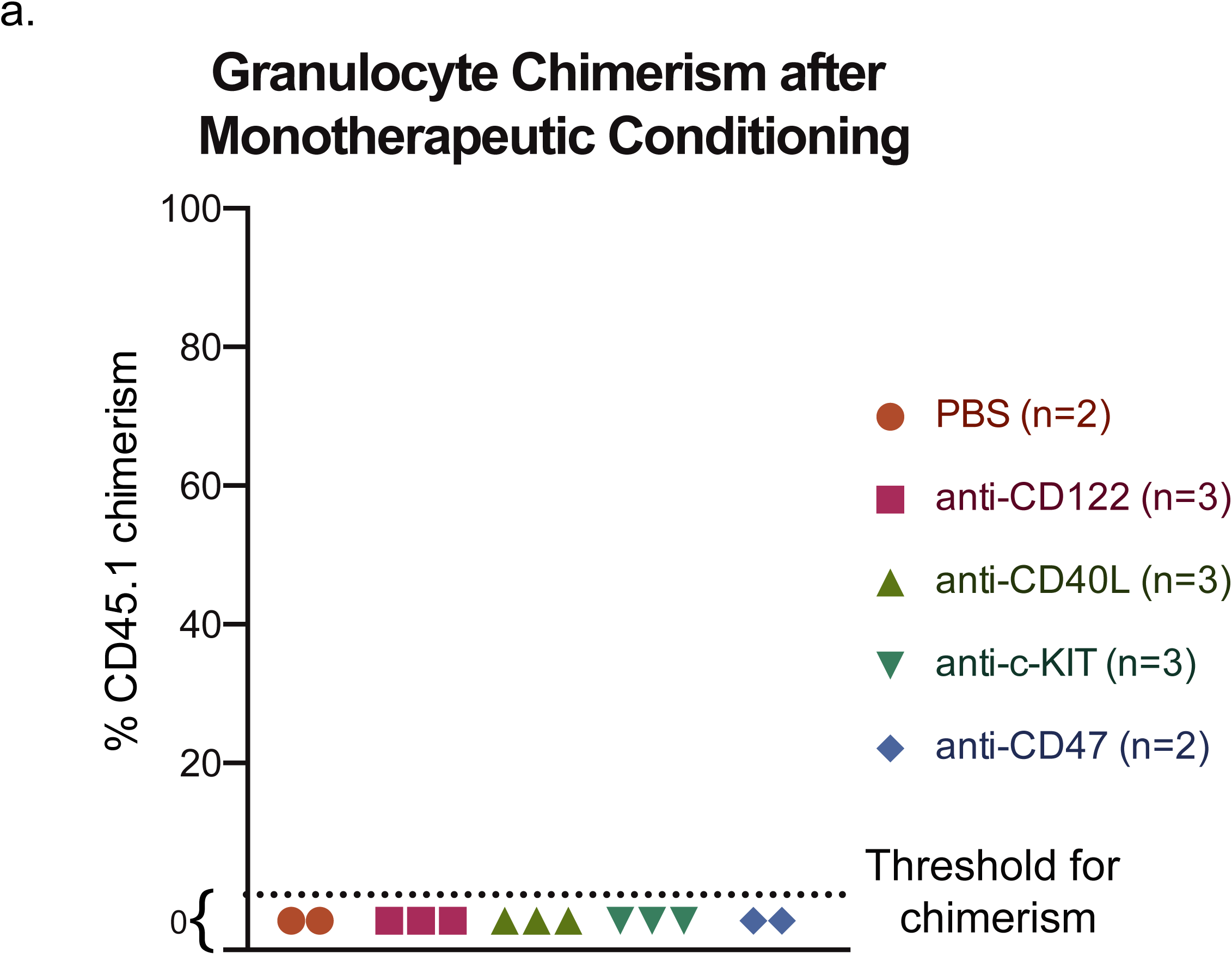
(a) Donor granulocyte chimerism following monotherapeutic conditioning using monoclonal antibodies was performed at 20 weeks (n=2-3).

**Extended Data Figure 3.**
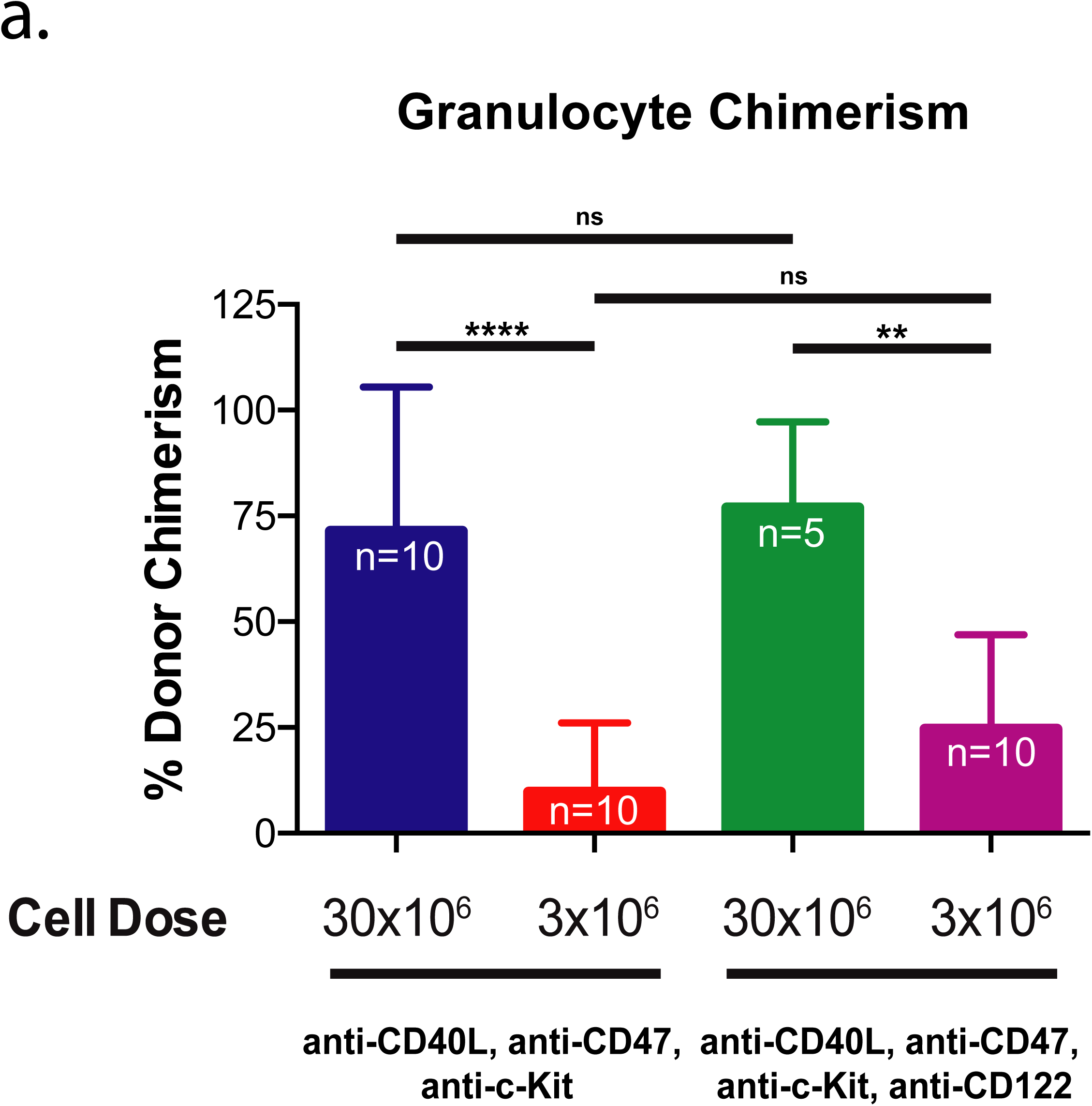
(a) 3-week granulocyte chimerism following variations in cell dose and use of anti-CD122 (n=5-10). Data and error bars in (a) represent means ± SD; one way ANOVA was performed where **P≤0.01 and ****P≤0.0001.

**Extended Data Figure 4.**
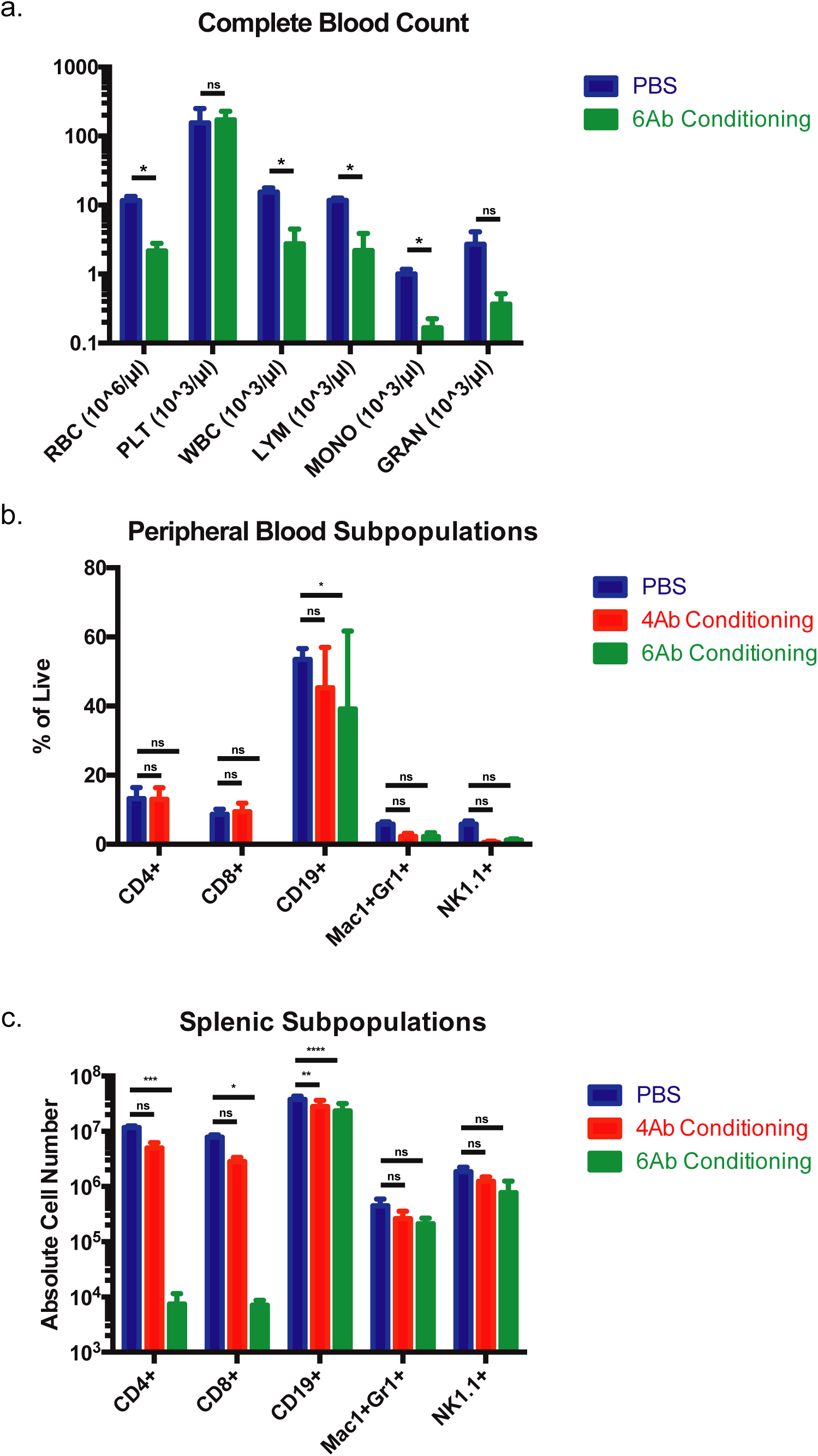
Complete blood count (a), peripheral blood subpopulations (b) and splenic subpopulations (c) from animals on Day 0 without transplantation (n=3). Data and error bars in (a-c) represent means ± SD; (a) multiple t-tests were performed where *P≤0.005, (b-c) two way ANOVA was performed where *P≤0.05, **P≤0.01, ***P≤0.001 and ****P≤0.0001.

**Extended Data Figure 5.**
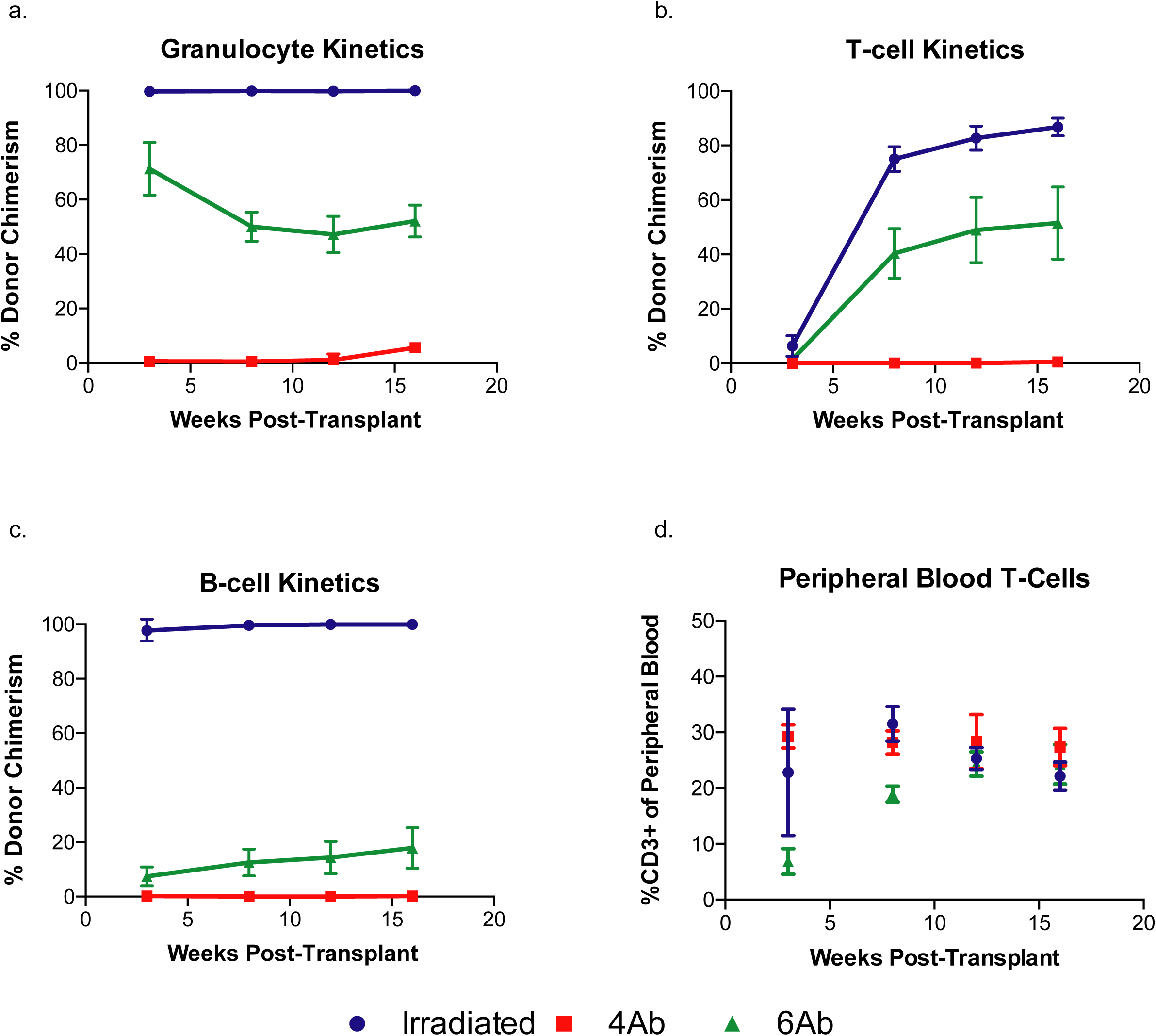
(a) Percentage of T-cells present in peripheral blood following transplant of LSK cells during 4Ab, 6Ab, and irradiation conditioning. Donor contribution following 4Ab, 6Ab and irradiation conditioning for (b) T-cells, (c) B-cells, and (d) granulocytes over 16 weeks.

**Extended Data Figure 6.**
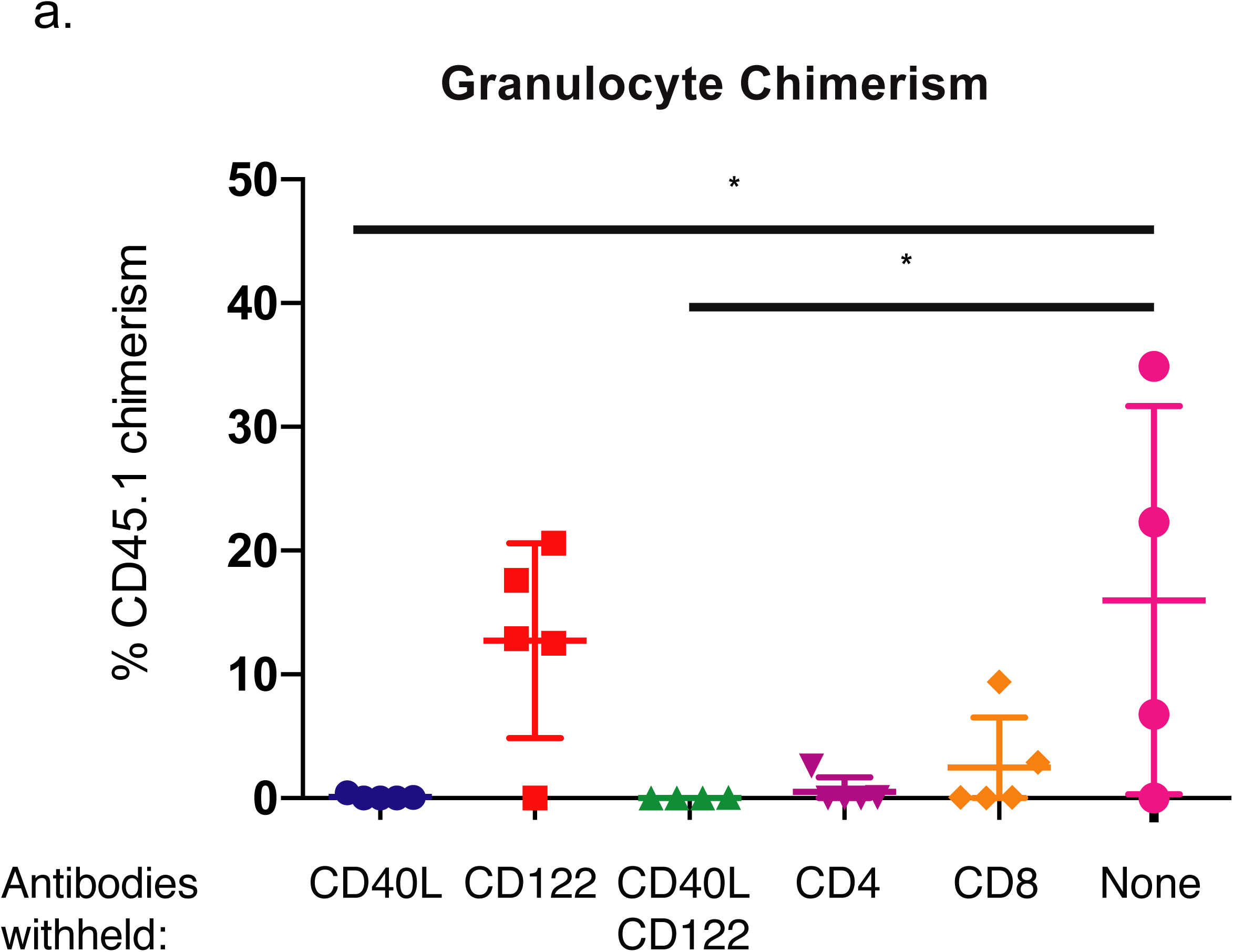
(a) 16-week granulocyte chimerism following variations of 6Ab conditioning. Data and error bars in (a) represent means ± SD; (a) one way ANOVA was performed where *P≤0.05.

Extended Data Video 1. Video of a representative HSC donor neonatal heart in the pinna of a HSC transplanted mouse. The viability and tolerance to these hearts was monitored by looking for beating of donor tissue, as shown in the video.

